# A Modular Approach to Active Focus Stabilization for Fluorescence Microscopy

**DOI:** 10.1101/2020.09.22.308197

**Authors:** Birthe van den Berg, Robin Van den Eynde, Baptiste Amouroux, Marcel Müller, Peter Dedecker, Wim Vandenberg

**Affiliations:** Department of Computer Science, KU Leuven, Belgium; Department of Chemistry, KU Leuven, Belgium; Faculty of Physics, Bielefeld University, Germany; University Paris-Saclay, France

## Abstract

Fluorescent time-lapse experiments often suffer from focus drift, regularly rendering long measurements partially unusable. Frequently, this instability can be traced back to the specific mechanical components of the setup, but even in highly robust implementations z-drift occurs due to small temperature fluctuations which are hard to avoid. To resolve this issue, microscope manufacturers often offer their own interpretation of out-of-focus correction modules for their flagship instruments. However, self-assembled or older systems typically have to fend for their own or adapt their measurements to circumvent drift effects. In this manuscript, we propose a cost-efficient z-drift detection- and correction system that, due to its modular design, can be attached to any fluorescence microscope with an actuated stage or objective, be it in a custom or commercial setup. The reason for this wide applicability is specific to the design, which has a straightforward alignment procedure and allows sharing optics with the fluorescent emission path. Our system employs an infrared (IR) laser that is passed through a double-hole mask to achieve two parallel beams which are made to reflect on the coverslip and subsequently detected on an industrial sCMOS camera. The relative position of these beams is then uniquely linked to the z-position of a microscope-mounted sample. The system was benchmarked by introducing temperature perturbations, where it was shown to achieve a stable focus, and by scanning different positions while simulating a perturbation in the z-position of the stage, where we show that a lost focus can be recovered within seconds.

## 1. Introduction

Keeping a sample in focus remains a challenging, yet essential, task in optical microscopy. Typically, this is most problematic in high-resolution imaging experiments where there is a limited depth of focus and a minimal discrepancy in the z-position can effectively render the collected data unusable. However, even low-resolution imaging can be impacted when time-lapse experiments are performed over longer time periods. Focus drift arises from a range of sources of which the most common ones are vibrations (e.g., fans), mechanical instabilities (e.g., loose gear), and thermal fluctuations (e.g., room temperature, laser heating up the sample). Although microscope designs have become increasingly more stable, and good practices have been developed, focus drift remains a practical, omnipresent and notable issue. Various techniques and systems have been proposed to detect and correct focus drift. While image-based detection methods [1, 2] rely on complex software that analyzes images and compares them with in-focus images captured beforehand, sensor-based detection methods have generally been more popular, using additional sensing circuits to detect drift and compensate for it using a control loop [3, 4].

Microscope manufacturers often build a z-drift compensation module into their instruments, such as Olympus’ TruFocus system [5], Leica’s Adaptive Focus Control (AFC) [6] and Nikon’s Perfect Focus System (PFS) [7]. These systems are rapidly becoming standard equipment on new motorized microscopes. However, as they are specific to the underlying microscope system, these commercial solutions may be more challenging to add to existing systems. For example, older systems, microscopes bought with a limited budget, and self-assembled systems that are often used in research environments are by default not equipped with an automated focus detection- and correction system. Several attempts have been made to control focus drift in those state-of-the-art microscope systems. However, most of these solutions suffer from ailments such as high cost, complex construction, lack of precision, slow detection and correction, and specificity to a particular context or technique [8–10].

In this manuscript, we present an automated focus detection- and correction system that is cost-efficient, easy to build and modular by design. Our setup is an optical z-drift detection- and correction system that uses an infrared light source split by a double-hole mask into two parallel beams. These beams are reflected on the interface between cover glass and sample, and captured by an industrial sCMOS camera. Software reads the data from the detector and computes the compensation necessary to maintain the focus position. The combination of the camera with the double-hole mask offers a smart approach for mitigating imperfections in the captured reflections [11], being less sensitive to flaws than quadrant detectors [9, 12] or line sensors [7].

Our automated z-drift detection- and correction system is implemented on a popular commercial microscope body (Olympus IX-71). However, due to its modularity and cost-effectiveness, it also opens up opportunities to mount it on other popular and conventional microscope systems. In particular, the system is optimized for microscopes in which the optical infrared path and the fluorescence path are shared, maximally eliminating reflections and noise. We show that our method works well through experiments with beads and a varying temperature, and on COS-7 cells with manual z-dispositioning.

## 2. Conceptual System Description

The idea of this work is to use a detection technique similar to that of Binh et al. [11] to automatically find the focus position for our microscope. This focus detection method uses a collimated infrared laser to determine the sample position. In front of the laser, we put a double-hole mask, which divides the collimated laser light into two parallel beams. These beams are sent through the microscope body onto the sample, which is carried by a glass coverslip. It is exactly this glass-sample interface that reflects the laser beams. The reflected beams are captured by an industrial sCMOS camera.

Figure 1 shows the possible ways for the laser beams to be reflected. The sCMOS camera detects two reflected laser spots. The horizontal distance between the detected spots is proportional to the z-position of the cover glass. The bigger the distance between the two detected spots, the further away is the sample from the zero of the z-position. Before starting an experiment, the user can define the perfect focus position of the sample on the microscope. The distance between the spots on that position determines the focusing distance. Control software then continuously adjusts the z-position to retain or regain focus, even when it is lost, by tuning the distance between the detected spots using a motorized stage or a mechanically actuated objective.

**Fig. 1.**
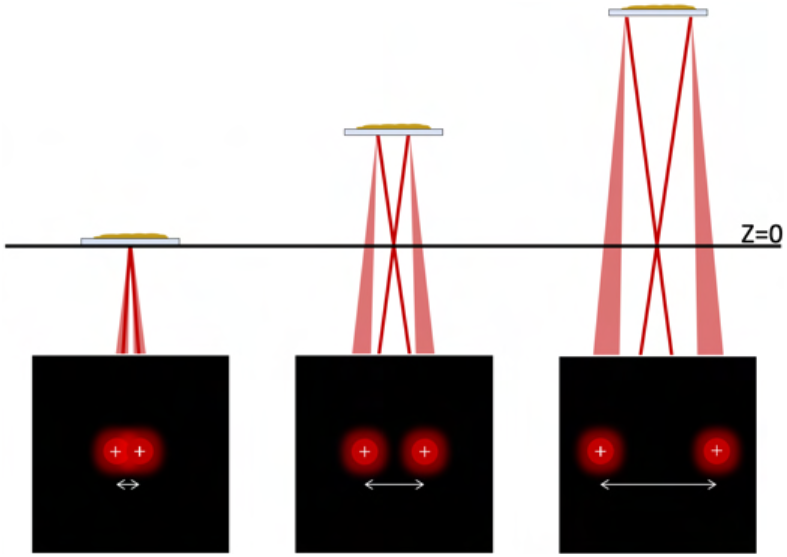
Correlation between z-position of sample and camera view.

The system has several advantages over existing approaches. Firstly, whereas other systems [7, 9, 12] typically have a dedicated path for the z-tracking laser, our system is designed to share their fluorescence path with the z-tracking laser path, which involves including dedicated components to filter out spurious reflections on intermediary components (e.g., tube lens). Secondly, for the implementation given in Appendix A, the total price of the system comes to less than €2000, a fraction of the cost of a commercial system. Thirdly, as the system focuses the two beams on the glass-sample interface, it presents a relatively safe way of using infrared light as any infrared light emanating from the objective is highly dispersed at eye height. Finally, the use of a two-spot design is beneficial. In techniques that employ a single back-reflected beam, the shape of the spot is of utmost importance, as in most cases a circular spot is assumed [9, 12–15] and any distortions of this beam profile directly result in a focusing error. To overcome this delicate issue, computing the centroid or center-of-mass was proposed [16]. However, even this algorithm is still influenced by imperfections at the border of the spot (Figure 2a). With a double-spot design, deviations are avoided, as long as the disturbance influences both beams in a similar way, leaving the distance between the two spots more or less the same (Figure 2b).

**Fig. 2.**
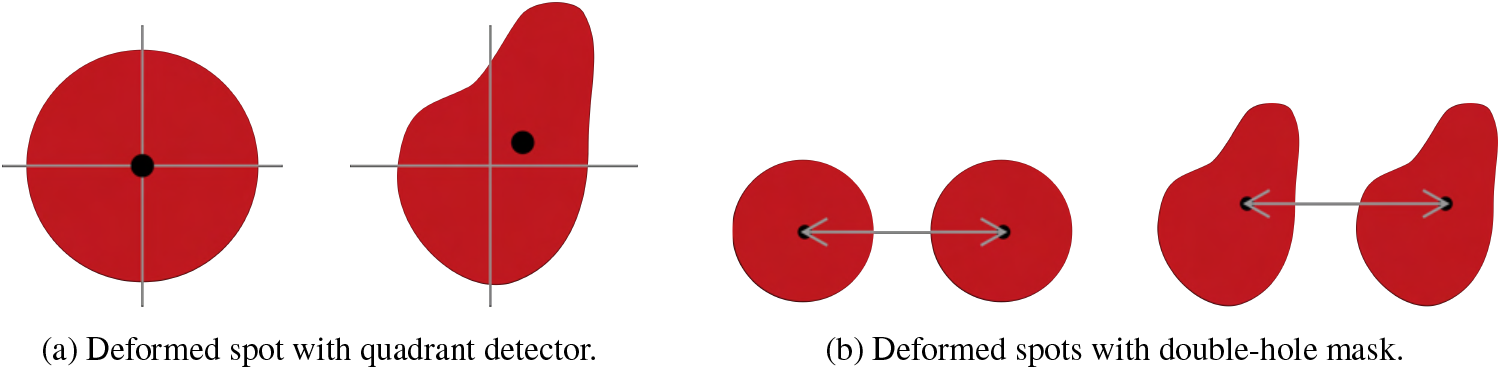
The influence of deformed spots on the focus for different detectors.

## 3. Practical System Description

### 3.1. Setup

Figure 3 gives a schematic overview of our automated focus detection- and correction system. The infrared light beam path is visualized in red, and the fluorescence path in blue. The detailed configuration is referenced in Appendix A.

**Fig. 3.**
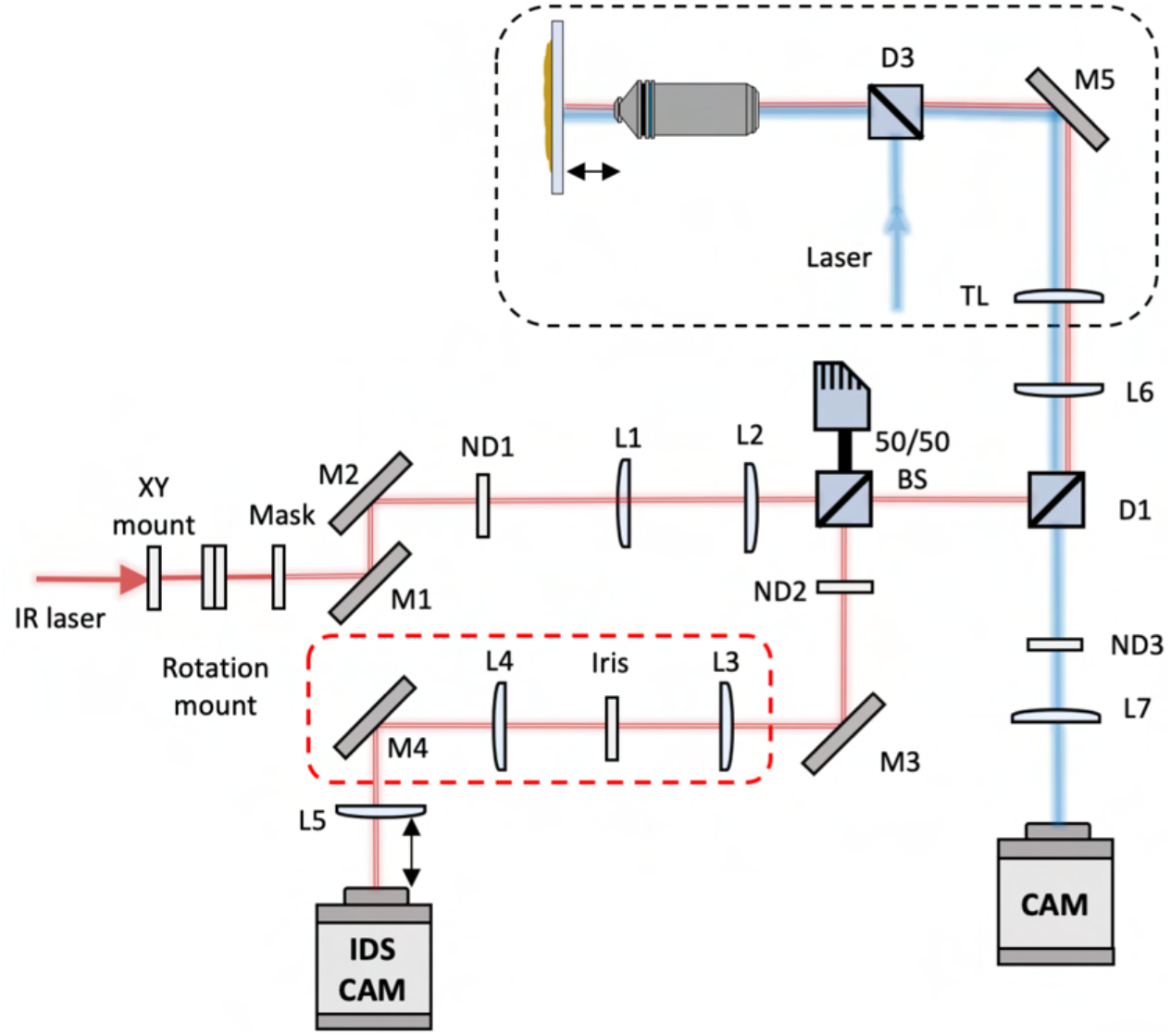
Schematic overview of the system. The blue line represents the fluorescence path, and the red line the path of the two parallel infrared beams. The beams are initiated at the IR laser (left).

#### Fluorescence Path

The sample is excited using a Spectra X light engine (Lumencor) (named “laser” in Figure 3). The excitation light is reflected by a dichroic mirror (D3) and passes the objective, illuminating the sample. The fluorescence path (blue) then initiates at the sample and travels through the objective and the dichroic mirror (D3), reflecting on the mirror M5 and passing the tube lens (TL) of the microscope body. In this setup, this part of the optical layout is implemented using a commercial olympus IX71 inverted fluorescence microscope, where in the scheme, the black dashed line indicates enclosure in the commercial microscope body. Beyond this commercial body, the fluorescence light travels further through a lens (L6), a dichroic filter (D1), an emission filter (ND3) and is finally focussed on a Hamamatsu ORCA-Flash4.0 LT+ Digital sCMOS camera using a lens (L7).

#### Infrared Beam Path

The infrared beam path starts from the infrared laser (the leftmost component in Figure 3). This laser is a 3.0 mW, 830 nm infrared laser, chosen to avoid overlap with the color channels used for fluorescence detection. This laser is mounted in a pitch/yaw adapter which, with addition of an XY-mount and a rotation mount, allows for full freedom in aligning the laser beam on a double-hole mask containing drilled holes that are 600 μm apart and 500 μm wide (Figure 4 shows the mask and its dimensions).

**Fig. 4.**
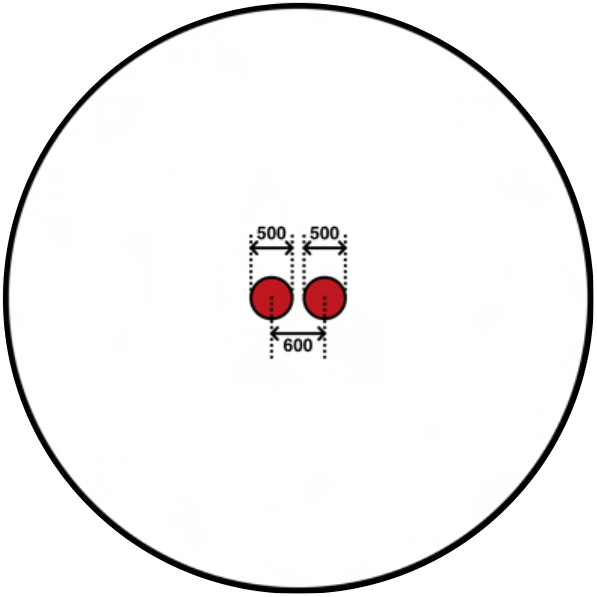
Cross-section of the double-hole mask with technical dimensions (μm).

Next, two mirrors M1 and M2 guide the two beams toward the objective-sample interface while a neutral density filter (ND1) reduces their power to less than 1 mW (safety category 2). Lenses L1 and L2 are set up as a 2F relay. A 50/50 beam splitter (BS) passes half the light coming from the laser into the direction of the microscope, while the reflected half is disposed in a beam dump. At this point, the tracking laser is reflected by D1 (separating the fluorescence and tracking path). After passing L6, the tube lens, and the objective, the two beams are reflected on the glass-sample interface and retrace their path up to the 50/50 beam splitter. There, the reflected light passes a laser-line clean-up filter (ND2) to remove any (fluorescent) background signal. Next, the light passes a spurious reflection filter (the red dashed frame in Figure 3). This extension filters out extra reflections that might disturb or overwhelm the spots to be measured. These reflections may originate from different sources, such as imperfections in optical components (beam splitter, dichroics), but in our case mostly consist of infrared light reflected from lenses shared with the fluorescence path. In this additional filter, mirror M3 is used to position the beams on the spatial filter (iris) to maximally filter out light noise, and lenses L3 and L4 are set up as a 2F relay. Finally, lens L5 focuses the two parallel infrared laser beams on the IDS camera sensor. The distance between this lens and the sensor chip of the IDS camera influences the shape of the spots and the distance between them. Figure 5 shows what the reflected laser beams look like when detected by the camera.

**Fig. 5.**
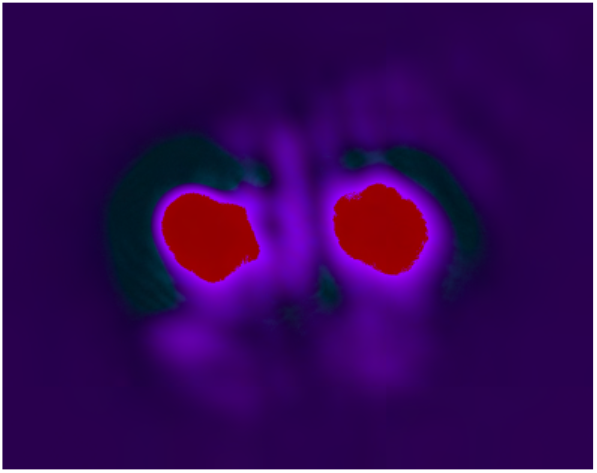
IDS camera view.

### 3.2. Software

The system is accompanied by supporting software, that determines the spot distances and adapts the z-position of the motorized stage when focus is lost. Briefly, as the IDS camera captures images, the software computes the center of the two spots using a center-of-mass algorithm based on the pixel intensities and calculates the distance between the centers of the two spots (using Figure 6). When, during an experiment, the automatic focus system is enabled and the calculated distance between the centers of the two spots changes, the algorithm sends an instruction to the actuated stage or objective to adapt its z-position proportional to the distortion of the computed distance. The software supports any motorized stage or objective positioner compatible with Micromanager [17, 18]. Currently, IDS cameras are used for spot detection but others can be added easily. Figure 7 shows the graphical user interface (GUI) of the software.

**Fig. 6.**
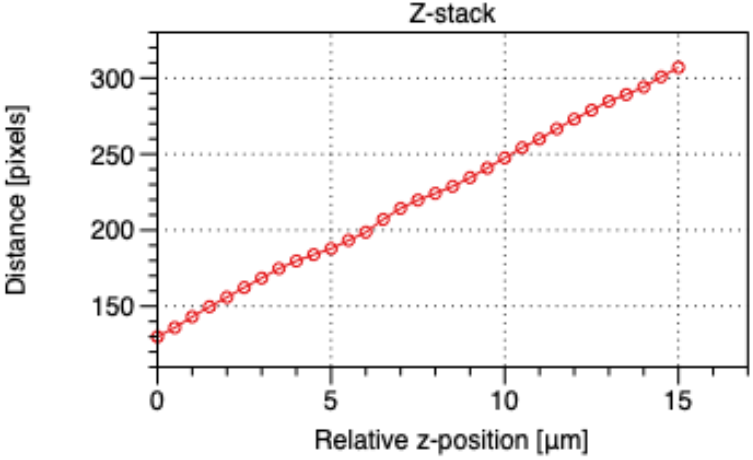
Z-stack of the autofocus system.

**Fig. 7.**
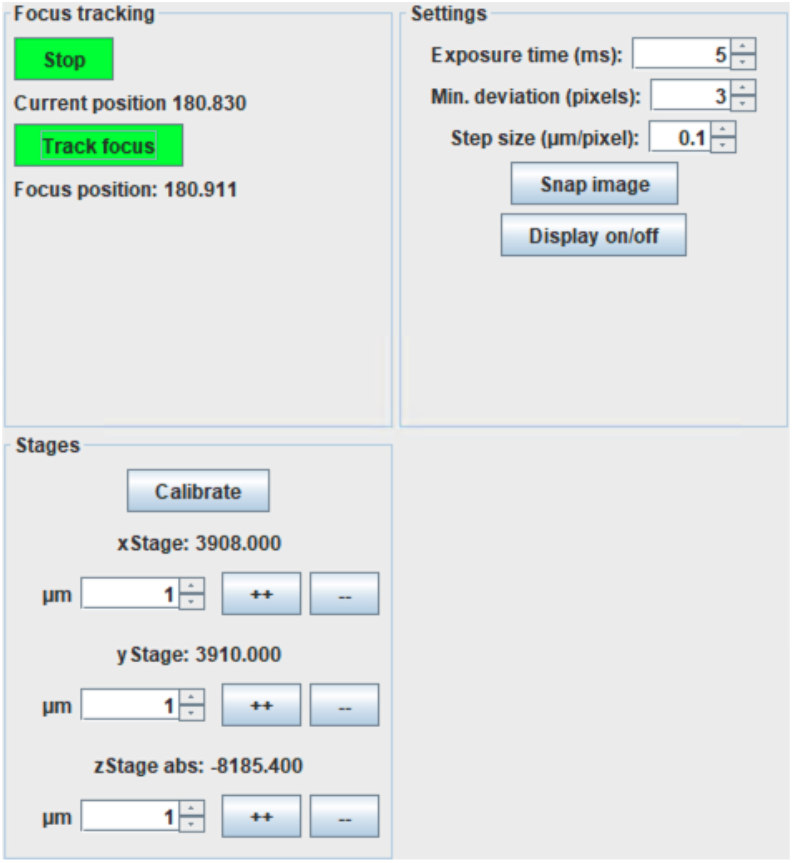
Graphical User Interface (GUI).

## 4. Experiments and Results

### 4.1. Stability Against Drift

We have validated our z-drift correction system by visualizing the mitigation of temperature-dependent drift on a fluorescence microscope. As a sample, we have used sparsely dispersed 200 nm Tetraspeck beads. We use an UMPlanFI 50× objective, with a numerical aperture of 0.80. The Spectra X light source is set to 50% at a band centered around 542 nm, a Chroma ZT561RDC dichroic filter(D3) and Chroma HQ572lp emission filter(ND3) are used. The exposure time is 0.1 s, and the maximal allowable deviation is set to 0.2 μm. In order to study Z-drift we use a time-lapse experiment, imaging every second, while the temperature is varied between 32°C and 45°C using the Okolab H301-T-UNIT-BL-PLUS stage top incubator. When the automated focus detection- and correction system is disabled, we clearly see the impact of the varying temperature on the z-position and the related focus of the sample: Figure 8 shows that the sample moves out of focus when the temperature is changing. The z-position and focusing distance change similarly and do not stay between the bounds (green and purple) that were set for the sample to be in focus.

**Fig. 8.**
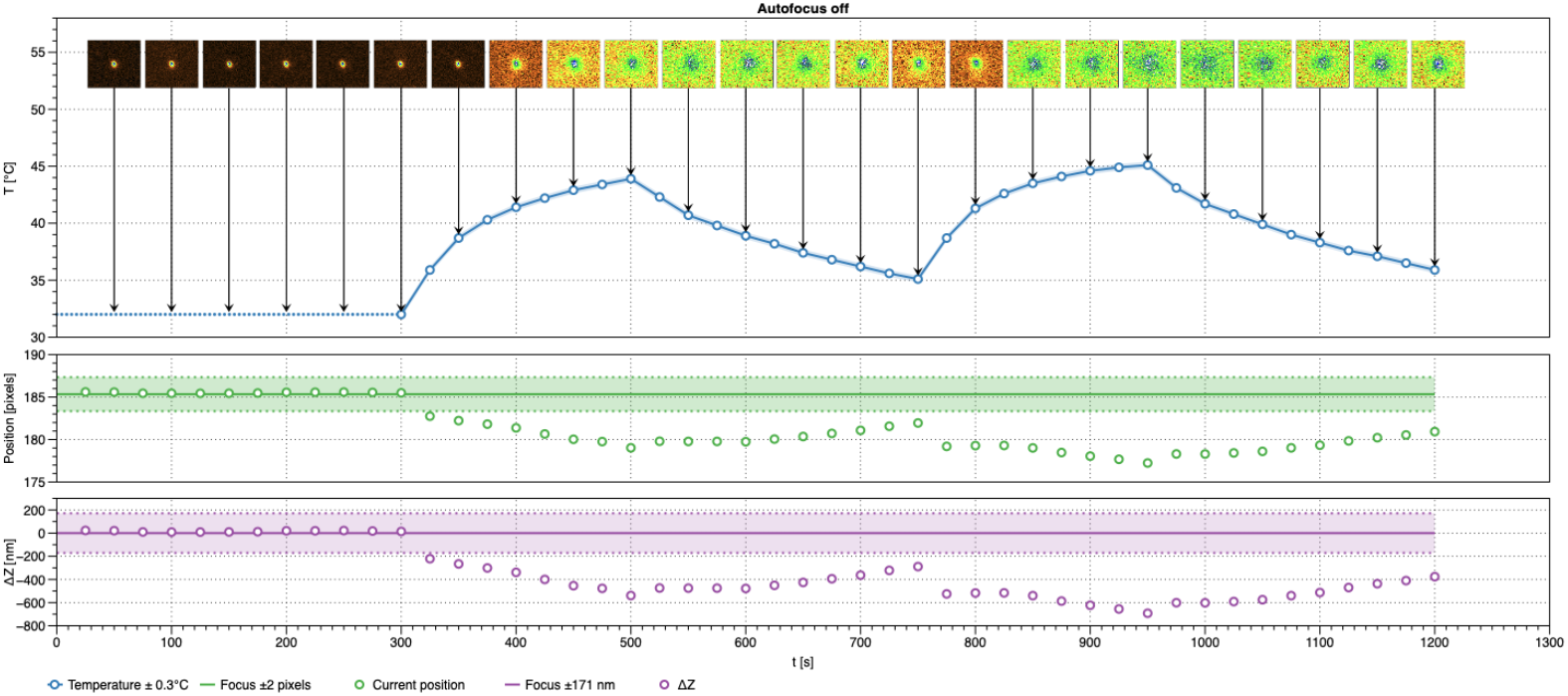
Time-lapse experiment with autofocus deactivated.

When we repeat the same experiment with the focus detection- and correction system enabled, Figure 9 shows that the images remain in focus. The focusing distance and z-position also stay between the predefined desired bounds. Similar results were achieved with a UPlanSApo 10× objective with a numerical aperture of 0.40 and a working distance of 3.1mm (data not shown).

**Fig. 9.**
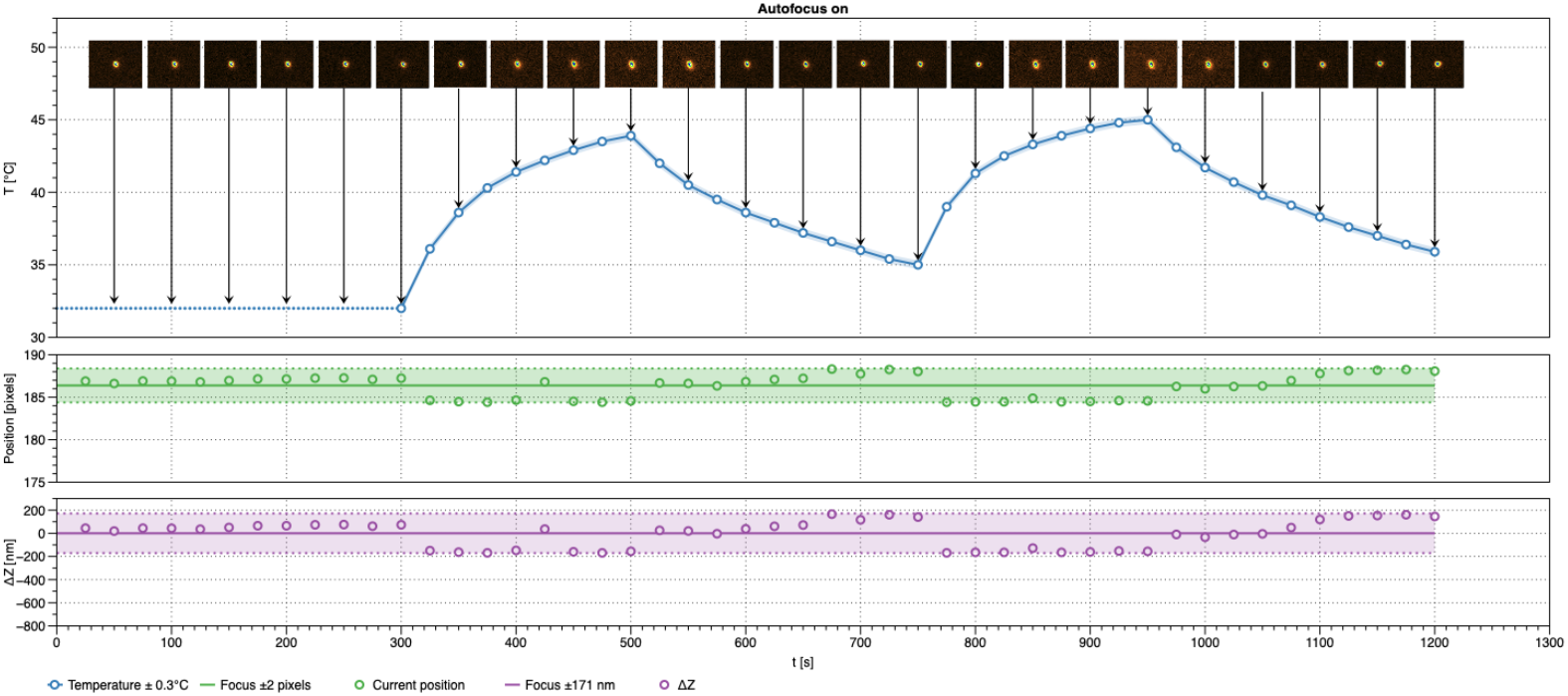
Time-lapse experiment with autofocus activated.

### 4.2. Stability Against Focus Perturbations

To study the stability of our system on focus perturbations, we image COS-7 cells expressing a fluorescent protein (detailed in Appendix B). We use an UMPlanFI 50× objective, with a numerical aperture of 0.80. During fluorescent imaging, the Spectra X light source is set to 10% at a band centered around 438 nm and a Chroma ZT561RDC dichroic filter (D3) and Chroma HQ572LP emission filter (ND1) are used. During focus tracking, the Chroma T455LP dichroic filter (D3) is used instead. The exposure time is set to 5s, and the maximal allowable deviation to 0.25 μm. Next, we loop over four distinct positions (Figure 11) and at each position we perturb the z-focus by a known amount. Then, we give the automatic focus detection- and correction system 20s to re-find focus. After these 20s, we change the dichroic filter (D3) and acquire an image of the cells. For each of the positions, we defined the ideal focusing distance before starting the experiment.

The results for one position are summarized in Figure 10 and Figure 12. The top row of Figure 10 shows the acquired images for different amounts of defocus with the focus detection- and correction system disabled. In the lower row, the automatic focus detection- and correction system is enabled and the sample nicely stays in focus, even when moving a 500 μm away from the desired focusing position.

**Fig. 10.**
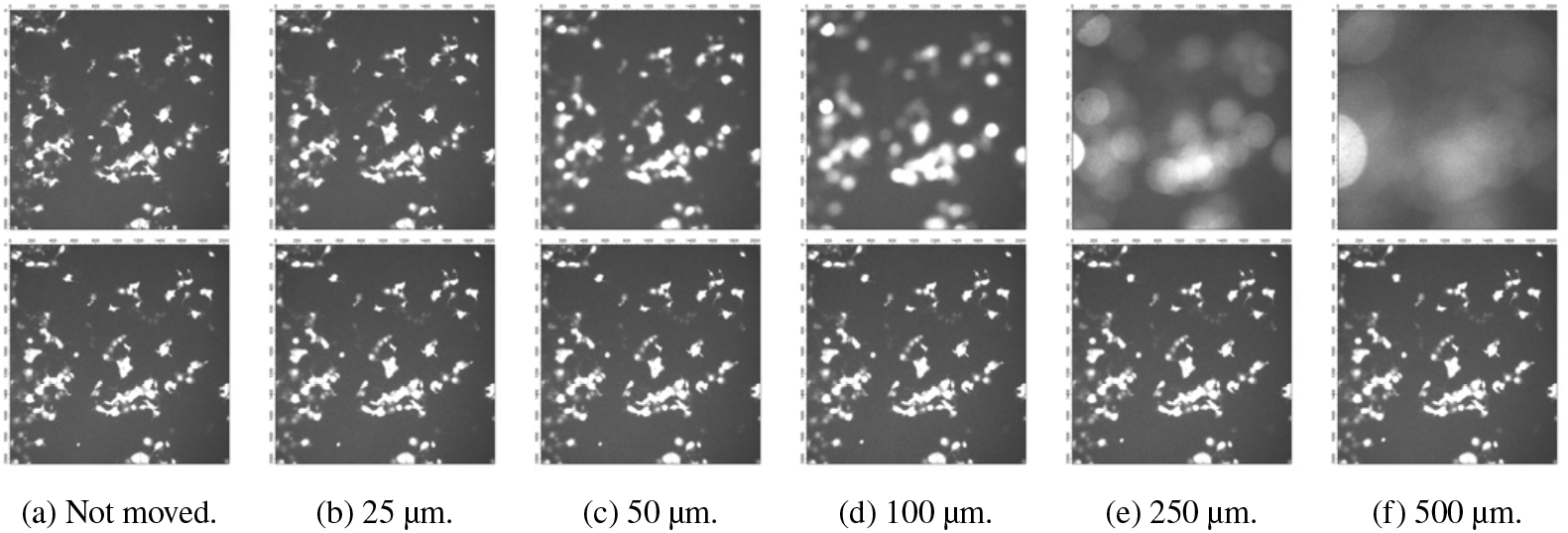
Acquired images for the experiment with cos7 cells for a single position. The first row shows the images when the automatic focusing system is off; in the second row it is on. In the respective images, we have moved down the z-position of the sample with the indicated amount.

**Fig. 11.**
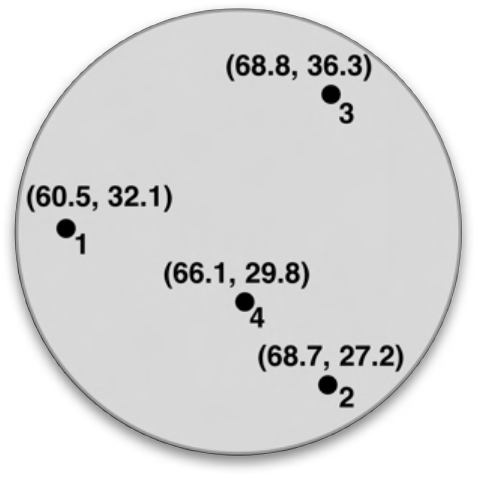
Four positions on the coverslip for imaging the COS-7 cells.

**Fig. 12.**
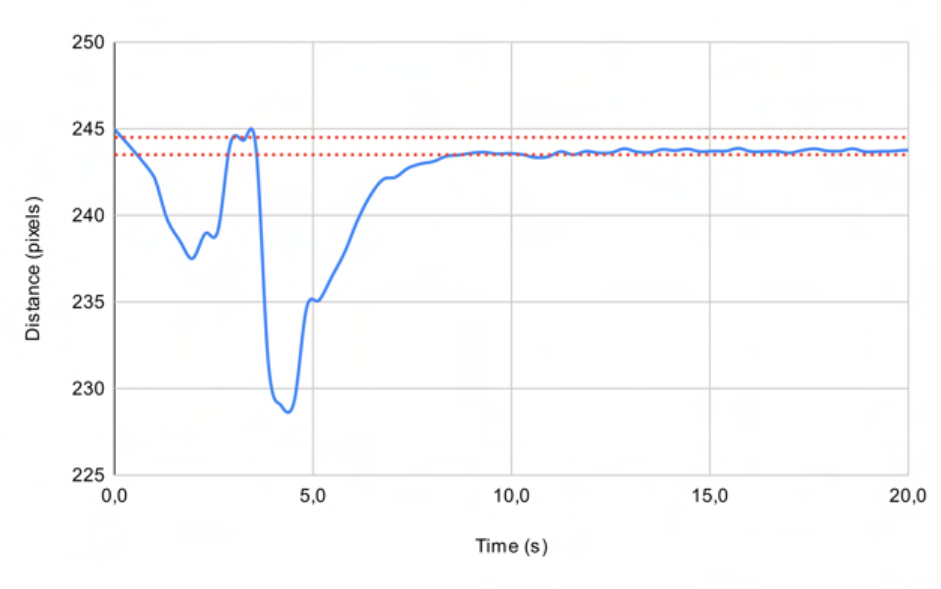
Focusing distance in function of time for position 1 (move 50 μm).

Figure 12 shows the distance measured between the detected spots from the infrared beams as a function of time after a perturbation in units of camera pixels (one camera pixel corresponds to 0.85 μm in the sample). The red dotted horizontal bars show the desired range for the distance between the spots for which the sample is considered to be in focus. The desired focusing distance is 244.0 pixels with a minimal deviation of 0.5 pixels. The blue line shows the movement of the spots when adapting to the desired focusing distance. The first local minimum in the curve is caused by the delay of the stage moving to the desired position. The second local minimum in the curve occurs when we perturb the z-focus (here by 50 μm) and give the autofocus system time to adapt. It finds back its desired focus within about 5 seconds.

## 5. Conclusion

We have described and built an automated out-of-focus detection- and correction system that is compatible with any microscope that has an actuated stage or objective. This automatic focusing system keeps a sample in focus during time-lapse experiments where z-drift occurs. The proposed system is cost-effective and robust, using a back-reflected infrared beam to detect the z-position of the stage or objective. The system is modular and can be easily extended with additional hardware components or added to a different microscope. In this case it is implemented as an aftermarket non-invasive add-on for a microscope where it shares the conventional detection path.

## Acknowledgements

Robin Van den Eynde thanks the FWO for a doctoral fellowship.

## Notes

### Competing Interest Statement

The authors have declared no competing interest.

### Summary of Updates

Novel experiments with cos7 cells, new title

## References

1. S. Yazdanfar, K. B. Kenny, K. Tasimi, A. D. Corwin, E. L. Dixon, and R. J. Filkins, “Simple and robust image-based autofocusing for digital microscopy,” Opt. Express 16, 8670–8677 (2008).

2. S. Li, X. Cui, and W. Huang, “High resolution autofocus for spatial temporal biomedical research,” The Rev. scientific instruments 84, 114302 (2013).

3. Y. Liron, Y. Paran, N. Zatorsky, B. Geiger, and Z. Kam, “Laser autofocusing system for high-resolution cell biological imaging,” J. microscopy 221, 145–51 (2006).

4. A. Pertsinidis, Y. Zhang, and S. Chu, “Subnanometre single-molecule localization, registration and distance measurements,” Nature 466, 647–51 (2010).

5. “TruFocus z drift compensator,” https://www.olympus-lifescience.com/en/microscopes/inverted/ix83/trufocus/!cms[focus]=cmsContent6373. Accessed: 2021-12-15.

6. “Leica adaptive focus control (afc),” http://major2020.syis.com.tw/webfiles/40a7c99d-a81e-4617-9fd4-87531a9b59c8.pdf. Accessed: 2021-12-15.

7. “The nikon perfect focus system (pfs),” https://www.microscopyu.com/tutorials/the-nikon-perfect-focus-system-pfs. Accessed: 2021-12-15.

8. W. Liu, Z. Zhong, and J. Ma, “Simple way to correct the drift in surface-coupled optical tweezers using the laser reflection pattern,” Opt. Express 29, 18769–18780 (2021).

9. M. Bathe-Peters, P. Annibale, and M. J. Lohse, “All-optical microscope autofocus based on an electrically tunable lens and a totally internally reflected ir laser,” Opt. Express 26, 2359–2368 (2018).

10. J. Luo, Y. Liang, and G. Yang, “Dynamic scan detection of focal spot on nonplanar surfaces: Theoretical analysis and realization,” Opt. Eng. - OPT ENG 50 (2011).

11. B. Cao, P. Hoang, S. Ahn, J.-O. Kim, H. Kang, and J. Noh, “In-situ real-time focus detection during laser processing using double-hole masks and advanced image sensor software,” Sensors 17 (2017).

12. W.-Y. Hsu, C.-S. Lee, P.-J. Chen, N.-T. Chen, F.-Z. Chen, Z.-R. Yu, C.-H. Kuo, and C.-H. Hwang, “Development of the fast astigmatic auto-focus microscope system,” Meas. Sci. Technol. 20, 045902 (2009).

13. C.-S. Liu, P.-H. Hu, Y.-H. Wang, S.-S. Ke, Y.-C. Lin, Y.-H. Chang, and J.-B. Horng, “Novel fast laser-based auto-focusing microscope,” in SENSORS, 2010 IEEE, (IEEE, 2010), pp. 481–485.

14. C.-S. Liu, P.-H. Hu, and Y.-C. Lin, “Design and experimental validation of novel optics-based autofocusing microscope,” Appl. Phys. B 109, 259–268 (2012).

15. C. Gu, H. Cheng, K. Wu, L.-J. Zhang, and X.-P. Guan, “A high precision laser-based autofocus method using biased image plane for microscopy,” J. Sensors 2018, 8542680:1–8542680:6 (2018).

16. C.-S. Liu, K.-W. Lin, and S.-H. Jiang, “Development of precise autofocusing microscope based on reduction of geometrical fluctuations,” in 2012 Proceedings of SICE Annual Conference (SICE), (IEEE, 2012), pp. 967–972.

17. A. D. Edelstein, M. A. Tsuchida, N. Amodaj, H. Pinkard, R. D. Vale, and N. Stuurman, “Advanced methods of microscope control using μ manager software.” J. biological methods 1 2 (2014).

18. A. D. Edelstein, N. Amodaj, K. Hoover, R. D. Vale, and N. Stuurman, “Computer control of microscopes using ţmanager,” Curr. Protoc. Mol. Biol. 92 (2010).

19. J. Fischer, D. Renn, F. Quitterer, A. K. Radhakrishnan, M. Liu, A. A. Makki, A. Rajjaka, S. A. Ghoprade, M. Rueping, T. Arold, M. Groll, and J. Eppinger, “A robust and versatile host protein for the design and evaluation of artificial metal centers,” ACS Catal. 9, 11371–11380 (2019).

