## Appendix A, Appendix B for "A Modular Approach to Active Focus Stabilization for Fluorescence Microscopy"

### A. List of Components

This section gives an overview of the necessary components to build the system, together with their reference number, their price and a description.

| # | Item | Part number | Unit price (€) | Description |
|---|---|---|---|---|
| 1x | Infrared laser | CPS830 | 103,28 | Collimated Laser Diode Module, 830 nm, 3.0 mW, Elliptical Beam, 11 mm Housing |
| 1x | Tilting mount | KAD11F | 63,44 | Threaded Kinematic Pitch/Yaw Adapter for 11 mm Cylindrical Components |
| 1x | Rotation mount | LRM1 | 84,86 | Rotation Mount for 1" (25.4 mm) Optics, External SM1 Threads |
| 1x | Double-hole mask | *Custom* | / | Space between holes: 600 µm, size of holes 500 µm |
| 3x | Mirror M1-M2-M3 | BB1-E03 | 70,31 | 1" Broadband Dielectric Mirror, 750 - 1100 nm |
| 2x | Lens L1-L2 | AC254-050-B | 85,83 | f = 50.0 mm, 1" Achromatic Doublet, ARC: 650 - 1050 nm |
| 1x | Beam splitter | BT610/M | 291,16 | Beam Trap, 400 nm - 2.5 µm, 30 W Max Avg. Power, Pulsed and CW, M4 Tap |
| 1x | Lens L3 | LA1509-B | 30,50 | N-BK7 Plano-Convex Lens, 1", f = 100 mm, AR Coating: 650 - 1050 nm |
| 1x | IDS camera | UI-3260CP Rev. 2 | 364,00 | Sony's 2.35 MP sensor IMX 249 (1936 x 1216 px) |
| 1x | Filter ND1 | NE10A-B | 72,06 | 25 mm AR-Coated Absorptive Neutral Density Filter, 650-1050 nm, SM1-Threaded Mount, OD:1.0 |
| 1x | Filter ND2 | FL830-10 | 92,46 | 1" Laser Line Filter, CWL = 830 ±2 nm, FWHM = 10 ±2 nm |
| 1x | Mirror M4 | BB1-E03 | 70,31 | 1" Broadband Dielectric Mirror, 750 - 1100 nm |
| 2x | Lens L4-L5 | AC254-050-B | 85,83 | f = 50.0 mm, 1" Achromatic Doublet, ARC: 650 - 1050 nm |
| 1x | Iris diaphragm | SM1D12SZ | 71,57 | SM1 Lever-Actuated Zero Aperture Iris Diaphragm, 11.9 mm Max Aperture |

The components needed for this system cost €1.797,89 in total, with as most expensive components the beam splitter and the IDS camera.

### B. Materials and methods

COS-7 cells were cultured in Dulbeccos modified Eagle medium (DMEM; Gibco, #31053028) supplemented with 10% (v/v) fetal bovine serum (FBS; Gibco, #70011036), 1× Glutamax (Gibco, #35050061) and 50 µg/ml gentamycin (GM; Sigma-Aldrich, #15750060) and maintained in a humidified incubator (37 °C, 5% $CO_2$). 24 h before transfection, cells were plated onto sterile 35 mm glass-bottom dishes (MatTek) and grown to 3070% confluence. Cells were subsequently transfected 48h before imaging using X-tremeGENE™HP DNA Transfection Reagent (Merck) and plasmid DNA, according to the manufacturers protocol. For optimal transfection results in each dish, 1 µg of plasmid DNA was introduced in 200 µL Opti-MEM™I Reduced Serum Medium (Optimem; Gibcon, #31985-062), and it was followed by the addition of 3 µL of X-tremeGENE™HP by pipetting thoroughly, then the mixture was left for at least 10 minutes incubation time at room temperature before dropwise addition to each dish. Cells were then grown for 48h before imaging. Before imaging dishes were washed 5 times with FluoroBrite DMEM (Gibco, #A1896702) supplemented with 10% (v/v) fetal bovine serum (FBS; Gibco, #70011036), 1× Glutamax (Gibco, #35050061) and 50 µg/ml gentamycin (GM; Sigma-Aldrich, #15750060) and let in 1mL of the supplemented FluoroBrite DMEM. Plasmid DNA used was available in the lab. It consists on a pcDNA3 construct with standard CMV promoter coding for the mTFP0.7 fluorescent protein [19].
